# A type-specific B cell epitope at the apex of Outer surface protein C (OspC) of the Lyme disease spirochete, *Borreliella burgdorferi*

**DOI:** 10.1101/2024.11.06.622094

**Authors:** David J. Vance, Grace Freeman-Gallant, Kathleen McCarthy, Carol Lyn Piazza, Yang Chen, Clint Vorauer, Beatrice Muriuki, Michael J. Rudolph, Lisa Cavacini, Miklos Guttman, Nicholas J. Mantis

## Abstract

Broadly protective immunity to the Lyme disease spirochete, *Borreliella burgdorferi*, is constrained by an overwhelming antibody response against type-specific epitopes on Outer surface protein C (OspC), a homodimeric helix-rich lipoprotein essential for early stages of spirochete dissemination in vertebrate hosts. However, the molecular basis for type-specific immunity has not been fully elucidated. In this report, we produced and characterized an OspC mouse monoclonal antibody, 8C1, that recognizes native and recombinant OspC type A (OspC_A_) but not OspC types B or K, and arrests *B. burgdorferi* motility independent of complement. Epitope mapping by HDX-MS localized 8C1’s epitope to a protruding ridge on the apex of OspC_A_ α-helix 3 (residues 130-150) previously known to be an immunodominant region of the molecule. Alanine scanning pinpointed 8C1’s core binding motif to a solvent exposed patch consisting of residues K_141_ H_142_ T_143_ D_144_. In parallel, analysis of 26 Lyme disease positive serum samples confirmed antibody reactivity with this region of OspC_A_, with residues E_140_ and D_144_ as being most consequential. Our results underscore the importance of α-helix 3 as a target of type-specific epitopes on OspC_A_ across mice and humans that should be taken into consideration in Lyme disease vaccine design.

## Introduction

Lyme borreliosis (or Lyme disease) is a potentially debilitating tick-borne infection caused by the spirochete *Borreliella burgdorferi* sensu latu (s.l). Following transmission via a tick bite, *B. burgdorferi* replicates locally in the skin, often presenting clinically as an expanding rash known as erythema migrans. If left untreated, the spirochete can disseminate to secondary tissues with possible neurologic and cardiac complications ^1,2^. In humans, *B. burgdorferi* infection is accompanied by a robust B cell response that has been associated with disease resolution ^3^. *B. burgdorferi*-specific antibodies also afford protection against reinfection, albeit with the caveat that immunity is restricted to strains expressing the same outer surface protein C (OspC) type ^4–11^.

OspC (BB_B19), a member of the small variable surface protein (Vsp) family unique to the *Borrelia* and *Borreliella* genera, is expressed by *B. burgdorferi* during tick-mediated transmission and in the early stages of mammalian infection, where it has an array of adhesin and immune evasion activities, including interactions with plasminogen and the complement component C4b ^12–19^. Structurally, OspC (∼23 kDa) is a helical-rich polypeptide that dimerizes to form a knob-shaped molecule anchored via a lipidated N-terminus to the spirochete’s outer membrane ^16^. In humans and other mammals, OspC is among the most immunoreactive of *B. burgdorferi*’s many outer surface proteins ^20^. It is also one of the most polymorphic of the spirochete’s many lipoproteins, with >30 known OspC types reported worldwide ^10,21–23^. Experimentally, OspC sequence diversity accounts for the limited cross protective immunity observed in serial challenge studies with *B. burgdorferi* expressing heterologous OspC types ^4,6,24–26^. Therefore, identifying the immunodominant and subdominant B cell epitopes on OspC is fundamental to understanding immunity to *B. burgdorferi* and has important implications for OspC-based vaccines ^27^.

## Results and Discussion

Monoclonal antibodies (mAbs) are powerful tools for identifying immunodominant and subdominant B cell epitopes on highly variable pathogen-associated surface antigens such as SARS-CoV-2 Spike, influenza virus haemagglutinin (HA), and HIV-1 envelope glycoprotein ^28–31^. The same approaches are being applied to OspC with the recent X-ray crystal structures of OspC bound to Fabs from mAbs B5 [PDB ID 7UIJ] and B11 [PDB ID 9BIF] ^32,33^. In pursuit of additional mAbs, we immunized groups of BALB/c mice with a mixture of recombinant OspC types A, B and K then screened splenic-derived B cell hybridoma supernatants for OspC-specific reactivity (see **Materials and Methods**). Supernatants from one particular hybridoma, 8C1, secreted IgG that recognized OspC_A_, but not OspC_B_ or OspC_K_ by Luminex (**S1 Fig**). Similarly, by dot blot, 8C1 hybridoma supernatants bound to *B. burgdorferi* strain B313 (OspC_A_) but not strains ZS7 (OspC_B_) or 297 (OspC_K_) (**S1 Fig**), providing further evidence that 8C1 recognizes an OspC_A_-restricted epitope.

To further characterize 8C1, the hybridoma was single cell cloned and the V_H_ and V_L_ coding regions were amplified from hybridoma-derived cDNA, subjected to DNA sequencing, and subsequently cloned as gBlocks™ into pcDNA3.1-based human IgG_1_ Fc and kappa light chain expression plasmids. Recombinant 8C1 IgG1 had an OspC reactivity profile identical to the hybridoma-derived 8C1, indicating successful reconstitution of the V_H_ and V_L_ pairing (**Fig 2; S2 Fig**). By flow cytometry, 8C1 bound to *B. burgdorferi* strain B31 (OspC_A_), but not a *B. burgdorferi ospC* deletion mutant or strains displaying OspC_B_ (ZS7) or OspC_K_ (297) (**Fig 1**). In addition, 8C1 induced both agglutination and alterations in outer membrane integrity of *B. burgdorferi* B313 (**Fig 1**), two functional activities shared with B5 and B11 that we have speculated may impact *B. burgdorferi* migration and transmission. The dissociation constant (K_D_) of 8C1 mAb for OspC_A_ was 11.2 nM (**S2 Fig**), while 8C1 Fab had a similar K_D_ for OspC_A_ of 10.7 nM (**S2 Fig**), as determined by biolayer interferometry (BLI).

**Fig 1.**
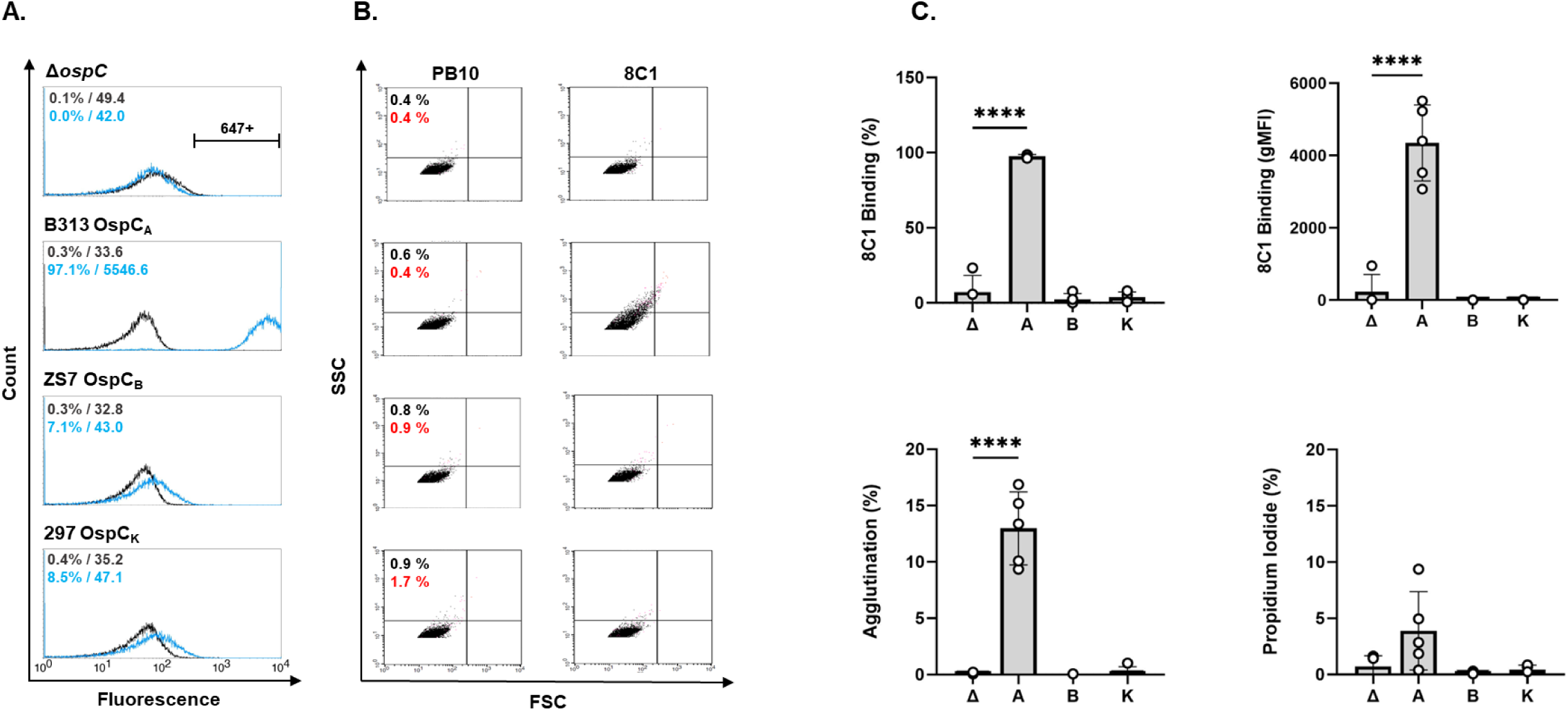
8C1 recognizes native OspC on the surface of *B. burgdorferi*. Cultures of B*. burgdorferi* B31A3, B313, ZS7 and 297 (as described in the text) were probed with 8C1 hIgG or an isotype control (PB10 hIgG), then stained with an Alexa Fluor 647-labeled anti-human IgG secondary antibody and subjected to flow cytometric analysis. (**A**) Representative histograms of 8C1 IgG (blue) and PB10 IgG (black) with text insets indicating % positive cells and gMFI for each strain examined; (**B**) Corresponding FSC (x-axis) and SSC (y-axis) dot plots representing event size and granularity, respectively. The percent of agglutinated events (black text insets) was calculated from the sum of UL+UR+LR quadrants, relative to the total events counted (20,000). The percent of events positive for propidium iodide (red text insets) are shown as red dots; (**C**) Combined flow cytometry analysis from n≥4 independent biological replicates with each symbol representing an independent experiment, the column representing the mean, and the error bars standard deviations. The OspC type (null, A, B and K) is plotted on the x-axis. The gMFI from PB10 was considered background was subtracted from values plotted. Asterisks specify statistical significance (p<0.0001; one-way ANOVA).

**Fig 2.**
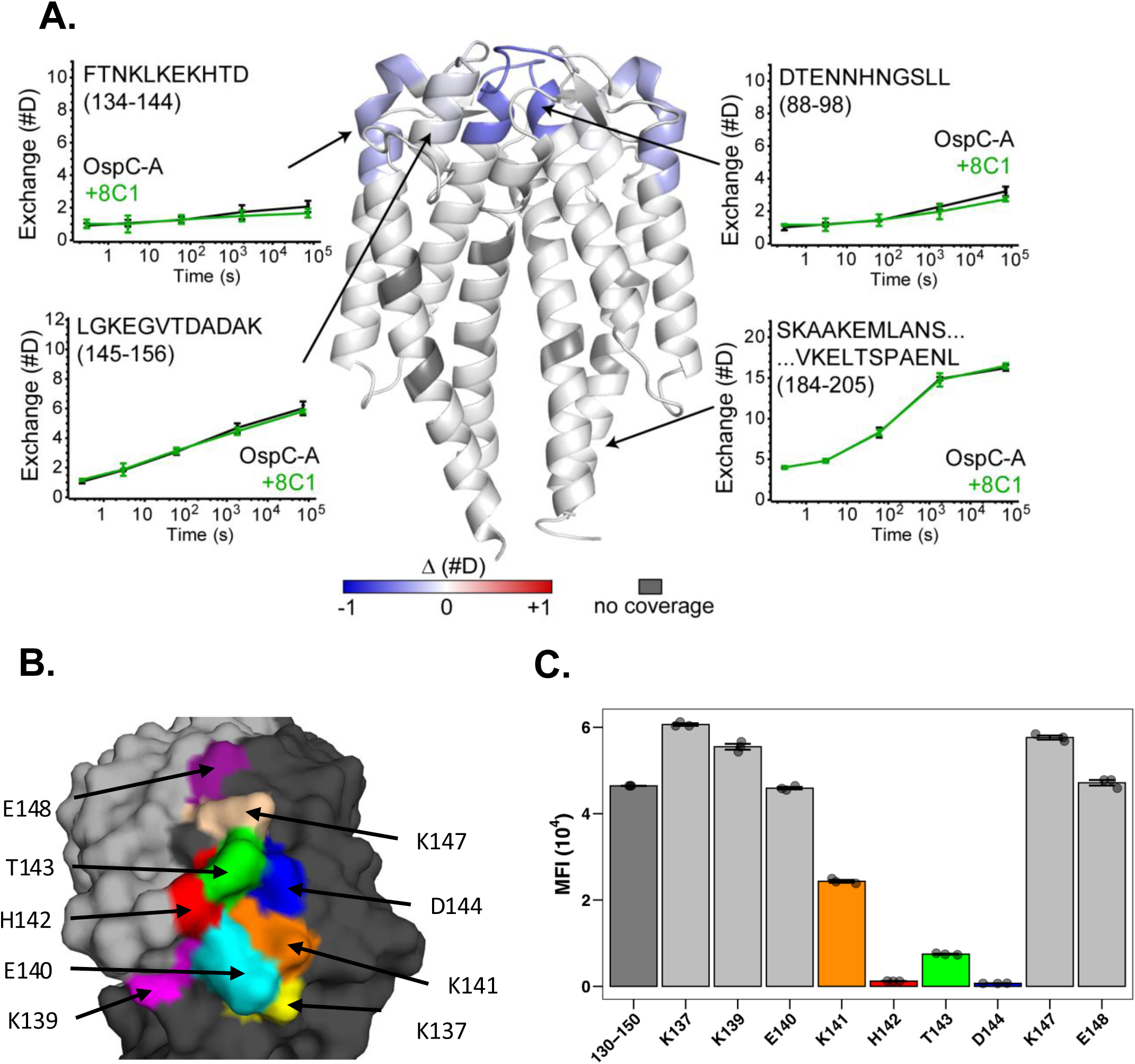
Localization of 8C1’s epitope to OspC_A_ residues 141-144. (**A**) ΔHDX in OspC_A_ upon incubation with MAb 8C1 are plotted on the structure of OspC_A_ [PDB ID 9BIF] in center of panel. Regions with increased protection are colored in blue and more exposed in red. Deuterium uptake plots for unbound OspC_A_ (black) and 8C1-OspC_A_ complex (green) are shown in plots for selected regions indicated by arrows. Symbols represent the mean ± SD from three independent measurements. (**B**) Surface representation of dimeric OspC_A_ [PDB ID 1GGQ; monomers colored gray and black] with 8C1’s putative epitope colored by residue; (**C**) 8C1 reactivity (MFI) with OspC_A_ peptide 130-150 (left column) or peptides 130-150 with Ala substitutions as indicated on the x-axis. The columns are color coded based on Panel B.

We next investigated whether 8C1 induces complement-dependent and/or - independent motility arrest of *B. burgdorferi*. Motility is critical for spirochete migration during tick transmission and within skin tissues ^34^. To circumvent issues associated with intrinsically low OspC expression by *B. burgdorferi* B31 in culture, we utilized a strain with an IPTG inducible *rpoS* allele, thereby activating native *ospC* expression *in trans* (see **Materials and Methods)**. In the absence of complement, *B. burgdorferi* cells treated with ≥10 µg/mL of 8C1 IgG were significantly less motile than isotype controls (**Table 1; S3 Fig**). The addition of 20% human complement further reduced motility of 8C1-treated cells, although the difference did not achieve statistical significance, indicating that the majority of 8C1’s effects on motility are complement-independent. This contrasts with mAb B5, which when evaluated in parallel had demonstrable complement-dependent borreliacidal activity (**Table 1; S3 Fig**).

**Table 1.**
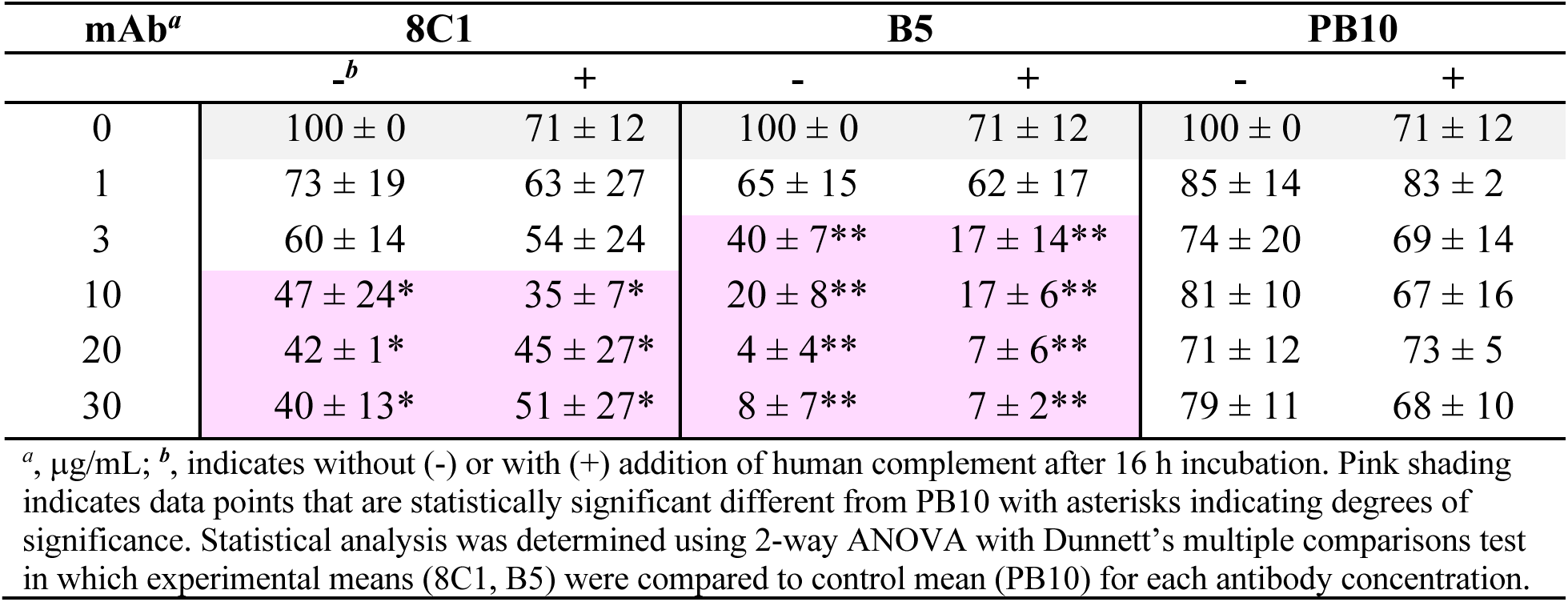
Antibody-mediated motility arrest (%) of *B. burgdorferi* GGW941.

| mAb<sup>a</sup> | 8C1 |  | B5 |  | PB10 |  |
| --- | --- | --- | --- | --- | --- | --- |
|  | -<sup>b</sup> | + | - | + | - | + |
| 0 | 100 ± 0 | 71 ± 12 | 100 ± 0 | 71 ± 12 | 100 ± 0 | 71 ± 12 |
| 1 | 73 ± 19 | 63 ± 27 | 65 ± 15 | 62 ± 17 | 85 ± 14 | 83 ± 2 |
| 3 | 60 ± 14 | 54 ± 24 | 40 ± 7** | 17 ± 14** | 74 ± 20 | 69 ± 14 |
| 10 | 47 ± 24* | 35 ± 7* | 20 ± 8** | 17 ± 6** | 81 ± 10 | 67 ± 16 |
| 20 | 42 ± 1* | 45 ± 27* | 4 ± 4** | 7 ± 6** | 71 ± 12 | 73 ± 5 |
| 30 | 40 ± 13* | 51 ± 27* | 8 ± 7** | 7 ± 2** | 79 ± 11 | 68 ± 10 |

<sup>a</sup>, $\mu\text{g/mL}$ ; <sup>b</sup>, indicates without (-) or with (+) addition of human complement after 16 h incubation. Pink shading indicates data points that are statistically significant different from PB10 with asterisks indicating degrees of significance. Statistical analysis was determined using 2-way ANOVA with Dunnett's multiple comparisons test in which experimental means (8C1, B5) were compared to control mean (PB10) for each antibody concentration.

We next sought to localize 8C1’s epitope on OspC_A_. In competitive binding assays, neither B5 nor B11 inhibited 8C1 from associating with OspC_A_ (data not shown), indicating that 8C1’s epitope is unlikely to be situated on the lateral face of OspC ^32,33^. Therefore, we turned to hydrogen-deuterium exchange mass spectrometry (HDX-MS) to identify regions of OspC that interact with 8C1 ^35^. A series of preliminary quench and digestion experiments revealed that proteolysis with Nepenthesin II without addition of urea generated the largest set of observable peptides for OspC_A_. After filtering out weak and overlapping signal there were 73 unique peptides remaining, resulting in a final sequence coverage of 98.8% with a redundancy of 5.3. The HDX-MS profiles of OspC_A_ without (unliganded) and with 2-fold molar excess of mAb 8C1 were compared. The magnitude of changes across OspC_A_ in the presence of 8C1 were minor, with only a few regions showing a statistically significant degree of protection. Among these few regions of OspC_A_ were peptides spanning residues 88-98 and 134-144, which correspond to the apex of the OspC_A_ dimer (**Fig 2; S1 File**). Minor protection was also detected along distal residues 145-156 but not the proximal peptide spanning residues 134-141. Based on this profile, we speculated that 8C1’s epitope is centered around OspC_A_ residues H_142_-T_143_-D_144_.

Residues 142-144 are nested within several previously reported mouse and human linear B cell epitopes, including the borreliacidal mAb, 16.22 (**Table 2**) ^24,36–38^. In fact, Marconi and colleagues refer to this region of OspC as loop 5 ^24,36^.

**Table 2.**
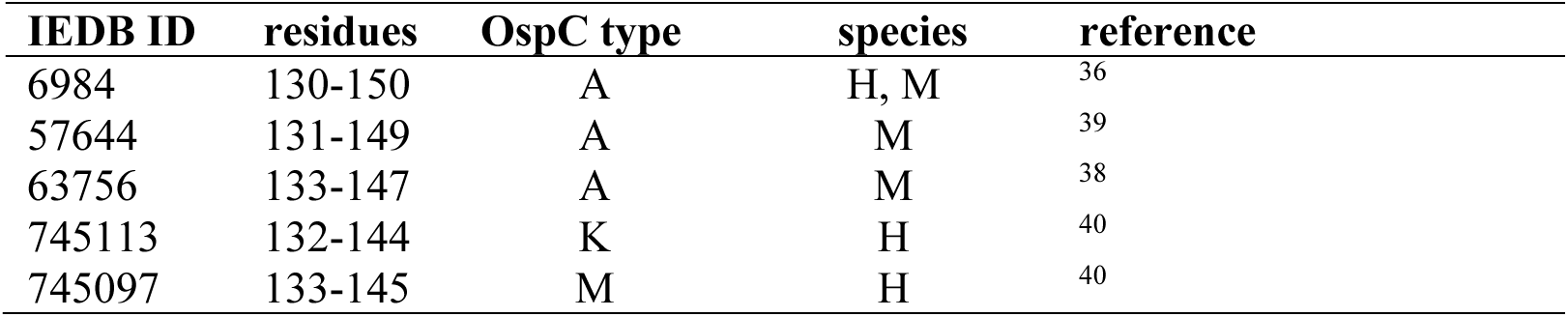
Reported OspC linear B cell epitopes overlapping with 8C1.

We therefore evaluated 8C1 reactivity with OspC-derived peptides spanning residues 130-150. In a microsphere immunoassay (MIA), 8C1 reacted strongly to OspC_A_ peptide 130-150 (**Fig 2C**) and to a lesser degree with peptide 132-146 (data not shown). By BLI, 8C1 Fabs had an affinity constant (K_D_) of 9.2 nM for peptide 130-150 compared to 10.7 nM for recombinant, dimeric OspC_A_, demonstrating that a linear epitope may account for a large proportion of 8C1’s binding energetics (**S4 Fig**). To define the critical residues associated with 8C1 interactions with the peptide, nine surface exposed residues within amino acids 130-150 (**Fig 2B**) were subjected to alanine (Ala) mutagenesis and the resulting biotinylated peptides were probed with 8C1. Ala substitutions at residues K_141_, H_142_, T_143_, and D_144_ each reduced 8C1 binding by 100- to 10,000-fold relative to native peptide 130-150 (**Fig 2**). These results are consistent with the HDX-MS analysis and demonstrate that K_141_, H_142_, T_143_, and D_144_ likely constitute important 8C1 contact points on OspC_A_. Moreover, those four residues alone may account for 8C1’s restricted OspC reactivity, as an alignment of 23 OspC types revealed that only Types A and C contain the K_141_ H_142_ T_143_ D_144_ motif (**Table 3**). In fact, the presence of an Ala at position 143 in 16 OspC types rather than OspC_A_ T_143_ is theoretically enough to completely abrogate 8C1 reactivity.

**Table 3.** Alignments of OspC residues 141-144.

| Type<sup>a</sup> | 141 | 142 | 143 | 144 | GenBank ID |
| --- | --- | --- | --- | --- | --- |
| A | K<sup>b</sup> | H | T | D | X69596 |
| A3 | E | Q | A | T | EF592541 |
| B | N | H | A | Q | CP001422 |
| C | K | H | T | D | DQ437462 |
| C3 | S | H | A | E | EF592543 |
| D | N | Q | A | E | CP001484 |
| D3 | S | H | A | V | EF592544 |
| E | E | H | A | V | AY275221 |
| E3 | S | H | G | E | EF592545 |
| F | G | N | A | Q | L42896 |
| F3 | E | H | T | D | EF592547 |
| G | S | N | A | D | AY275223 |
| H | E | H | A | S | CP001271 |
| H3 | S | H | G | N | FJ932733 |
| I | E | H | T | D | AY275219 |
| I3 | G | N | A | Q | FJ932734 |
| J | S | H | A | E | CP001535 |
| K | E | H | A | Q | AY275214 |
| L | E | N | V | A | EU375832 |
| M | S | H | A | E | CP001550 |
| N | S | H | A | Q | EU377775 |
| T | G | H | A | E | AY275222 |
| U | S | H | A | D | CP001493 |

<sup>a</sup>, OspC types derived from; <sup>b</sup>, Letters in red indicate positions of identity to the sequence of type A.

We recently extended the work by Buckles and colleagues that OspC types A, B and K-derived peptides encompassing residues 132-146 are recognized by sera from individuals positive for Lyme disease ^36,37^. Therefore, to investigate the contribution of the K_141_, H_142_, T_143_, and D_144_ motif in the peptide recognition in humans, OspC_A_ peptide 130-150 and the Ala mutants were probed with human Lyme disease serum samples (n=26 total) ^37^. Recognition of virtually every human serum sample was negatively affected by Ala substitutions at residues E_140_ or D_144_ (**Fig 3**). Ala substitutions elsewhere in the peptide enhanced antibody binding, a phenomenon observed by others for reasons possibly relating to epitope unmasking ^41,42^. These findings confirm that this region is highly immunogenic in lyme disease patients and suggests that there are human linear epitopes dependent on E_140_ and D_144_.

**Fig 3.**
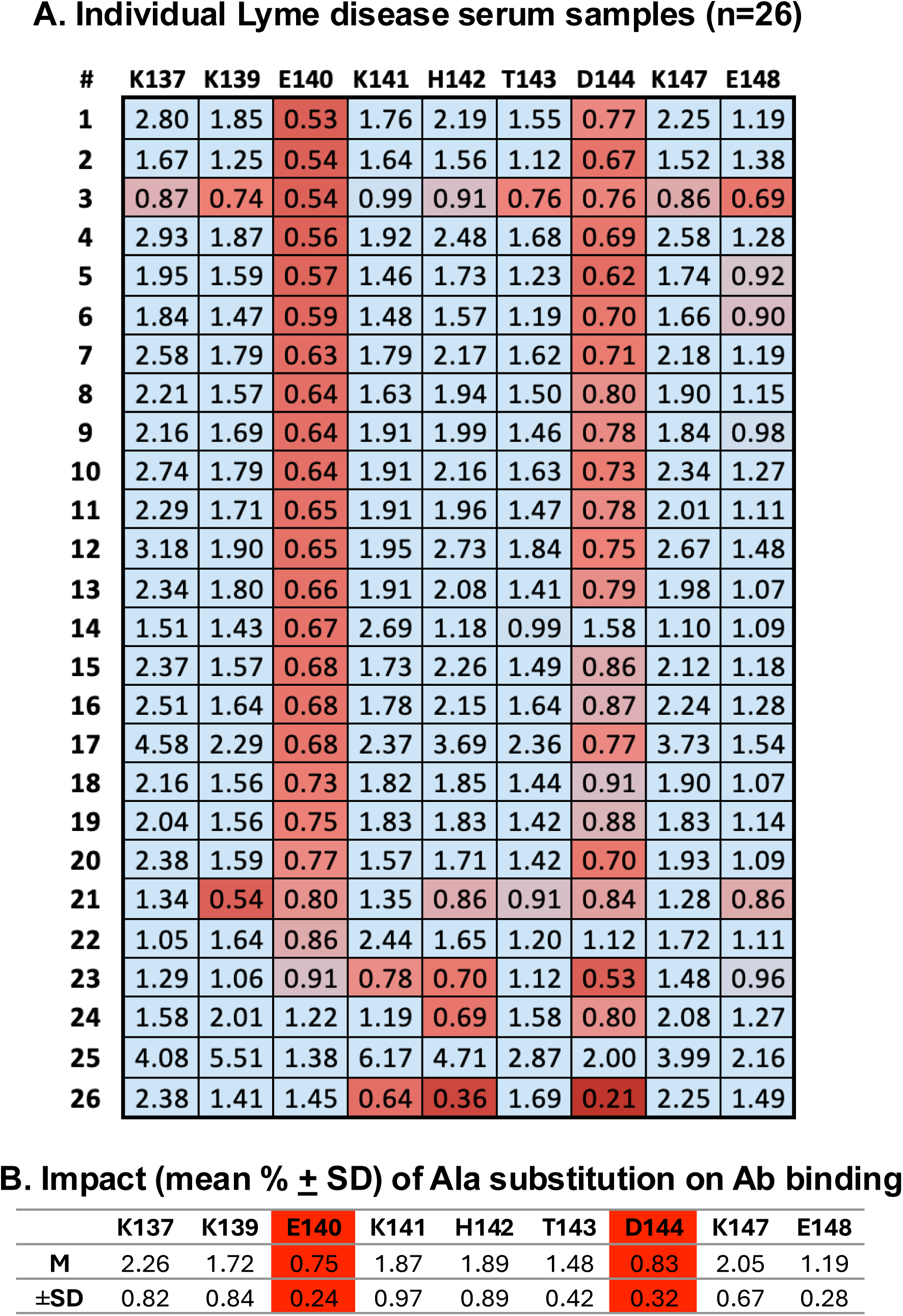
Recognition of OspC_A_ residues 137-150 by *B. burgdorferi* seropositive serum samples. (A) Lyme disease positive human serum samples (n = 26) previously determined to react with OspC_A_ residues 130-150 were subjected to MIA using 130-150 peptides with indicated single Ala substitutions, as decribed in the Material and Methods. The value in each box represents percent binding relative to the wild type peptide. Values <1 (colored red) indicate a reduction in peptide recognition. (B) Mean ± standard deviation of MFI ratios for each Ala substitution for all 26 serum samples.

## Conclusions

OspC defines strain-specific immunity to *B. burgdorferi* and therefore limits its utility as a vaccine ^4^. However, the B cell epitopes that constrain antibody reactivity to one OspC type at the expense of others are not fully characterized. In this report, we defined a linear epitope situated at the apex of OspC_A_ necessary for recognition by the mAb 8C1. The core of 8C1’s epitope consists of residues K_141_ H_142_ T_143_ D_144_, which constitutes a sequence variable but structurally conserved region on α-helix 3. Indeed, in our effort to better define the B cell epitopes on OspC, we inadvertently “rediscovered” an epitope hotspot ^24,38,40^. For example, Yang and colleagues identified this region as the target of the complement-independent, borreliacidal mouse mAb 16.22 ^38^. Peptide reactivity profiles led those investigators to conclude that 16.22 recognizes the core K_139_-E_140_-K_141_ motif. However, based on their results, it is equally plausible that 16.22 recognizes the KHTD motif just like 8C1. Either way, 16.22 and 8C1 most certainly have overlapping binding sites on OspC_A_ and share the capacity to induce *B. burgdorferi* motility and growth arrest.

Even before 16.22 was reported, Marconi and colleagues had identified residues 135-150 as being an immunodominant element on OspC, a region they referred to as loop 5, in mice and humans ^24,36^. Indeed, they made the case that loop 5 is highly conserved within a given OspC type but variable across types. For example, 53 of 57 (>90%) OspC_A_ sequences were identical across this region. Our results are consistent with that observation, with human immune sera being particularly prone to interactions with E_140_ and D_144_. More recently, Tokarz and colleagues demonstrated that antibody reactivity to analogous regions on OspC_K_ (residues 132-144) and OspC_M_ (residues 133-145) are diagnostic for Lyme disease ^40^. Thus, our report underscores the importance of the so-called loop 5 as a location of immunodominant and type-restricted epitopes on OspC in mice and humans with implications for both Lyme diagnostics and vaccine design.

## Materials and Methods

### OspC protein expression

Recombinant *B. burgdorferi* OspC_A_ (residues 38 to 201; PDB ID 1GGQ; UniProt ID Q07337) ^43^, OspC_B_ (residues 38 to 202; *B. burgdorferi* strain ZS7; PDB ID 7UJ2) ^44^ and OspC_K_ (residues 38 to 202; *B. burgdorferi* strain 297; PDB ID 7UJ6) ^45^ were expressed in *E. coli* strain BL21 (DE3) and purified by nickel-affinity and size-exclusion chromatography, as described ^32^.

### Bacterial strains and culture conditions

*B. burgdorferi* strains expressing OspC types A (B313), B (ZS7), K (297), and the *ospC* deletion strain B31A3ΔospC were cultured in BSK-II medium at 37°C with 5% CO_2_ to mid-log phase ^46^, collected by centrifugation (3,300 × *g*), washed, suspended in BSK II with 20% glycerol and stored at −20 °C until needed. Strain B31 (Cat# 35210) was obtained from the American Type Culture Collection (ATCC; Manasas,VA). Base BSK-II medium was prepared by the Wadsworth Center’s Tissue and Media Core Facility and filter sterilized (0.2 μm) prior to use. Cultures were maintained at 37°C with 5% CO_2_ and passaged by dilution (1:10,000) into fresh BSK-II medium. *B. burgdorferi* cultures were routinely inspected for culture viability and motility during *in vitro* culture maintenance prior to the initiation of any experiments.

### Dot blot

Bacterial strains expressing OspC types A (B313), B (ZS7), K (297), and deletion strain B31A3ΔospC were cultured in BSK-II medium at 37°C with 5% CO_2_ to mid-log phase, collected by centrifugation (3,300 × *g*), washed with PBS, and stored at −20 °C until needed. The bacterial pellets, and aliquots of recombinant OspC types A, B, and K, were diluted 10-fold in PBS before spotting on nitrocellulose membrane. PBS and an unrelated protein, rOspA, were included as negative controls. Incubation with 8C1 hybridoma supernatant, and processing and analysis of the dot blot were performed as described ^47^.

### Surface labeling and antibody-mediated agglutination of *B. burgdorferi*

*B. burgdorferi* strains expressing OspC types A (B313), B (ZS7), and K (297) were treated with 8C1 human IgG1 (10 µg/mL) and analyzed by flow cytometry, as recently described ^32^. Briefly, strains were cultured in BSK-II media minus gelatin at 33°C with 2% CO_2_ to mid-log phase, subcultured 1:10 into fresh medium medium, then grown at 23°C to early log phase. Cells were collected by centrifugation (3300 × *g*), washed with PBS, resuspended in media minus phenol red, and incubated at room temperature for 30 min. A total of 5 x 10^6^ cells in 50 µL volume were incubated with 8C1. Surface-bound antibodies, the cells were detected with goat anti-human IgG [H+L] cross-adsorbed secondary antibody Alexa Fluor 647 (Invitrogen). propidium iodide (0.75 µM; Sigma) was added to the culture for detection of membrane permeability. Analysis was conducted on a BD FACSCalibur flow cytometer (BD Biosciences Franklin Lakes, NJ). Cells were gated on forward scatter and side scatter to assess aggregate size and granularity. Alexa Fluor 647 fluorescence, PI staining, and agglutination were measured on 20,000 events using CellQuest Pro software (BD Biosciences, Franklin Lakes, NJ). Background reactivity of the PBS-immunized mouse serum for fluorescence and PI staining were considered baseline. Agglutination was calculated by the sum of events in the upper-left, upper-right, and lower-right quadrants relative to total events counted, as reported ^48^.

### Mouse immunizations and B cell hybridoma production

Animal studies were approved by the Wadsworth Center’s Institutional Animal Care and Use Committee (IACUC). Female BALB/c mice of ∼10 weeks of age were immunized with a combination of OspC Types A, B and K (20 μg each; 60 μg total) in 100 μL via the intraperitoneal route. The proteins were emulsified in 50% TiterMax Gold adjuvant TiterMax Gold (Sigma Aldrich, St Louis, MO). Mice were immunized three times at 3-week intervals, and OspC-specific titers were confirmed by indirect ELISA. Mice were boosted with OspC (without adjuvant) 4 days before being sacrificed.

Splenocytes were mixed 1:2 with Sp2/0 mouse myeloma cells and fused with PEG and subject to hypoxanthine-aminopterin-thymidine (HAT) selection ^49,50^. Supernatants were tested by multiplexed immunoassay (MIA) for reactivity with OspC types A, B and K (see below). Positive B cell hybridomas wells were cloned by two rounds of single cell dilution then and transitioned into hypoxanthine-thymidine (HT) medium.

### Direct and capture ELISAs

OspC (1 μg/mL) was coated on 96 well Immunoplate (Thermo Scientific, Waltham, MA) at a concentration of 1 μg/mL overnight at 4°C. Wells were blocked by 2% goat serum in 0.1% v/v PBS-Tween20 (PBST) for 2 h at room temperature. Two-fold serial dilutions of mouse serum starting from 1:100 were prepared in a separate dilution plate, and then transferred to the OspC coated plate for 1 h. Wells were washed with 0.1% PBST and anti-mouse IgG-HRP secondary antibody was added to detect bound mouse IgG. Wells were washed with PBST and then developed with SureBlue TMB (Seracare, Milford, MA). The reaction was quenched with 1M phosphoric acid, and the absorbance values were recorded at 450 nm by a SpectraMax iD5 spectrophotometer using SoftMax Pro 7 software (Molecular Devices, San Jose, CA)

### 8C1 V_H_ and V_L_ sequence determinations and IgG expression

The murine V_H_ and V_L_ cDNA sequences were determined as described ^51^. Briefly, total RNA was extracted from the 8C1 hybridomas via the RNeasy Mini Kit (Qiagen), reverse transcribed into cDNA using the Smartscribe Reverse Transcriptase Kit (Takara Bio, San Jose, CA) then used as template for PCR with reverse primers for the constant regions of the heavy, light kappa, and light lambda genes, as well as the universal forward primer added during reverse transcription. The PCR products were cloned into the Zero Blunt TOPO vector (Thermo Fisher Scientific) and transformed into Top Ten *E. coli* (Thermo Fisher Scientific). Several individual colonies from each transformation were picked, grown overnight, miniprepped to obtain the plasmid, and submitted for Sanger sequencing using the forward and reverse primers supplied in the Zero Blunt cloning kit. The the mouse immunoglobulin heavy- and light-chain genes (VH/VL) were cloned into pcDNA3.1 in-frame with human IgG1 and human kappa chain backbone (Genscript, New Jersey). Equal amounts of heavy- and light-chain plasmids were transfected into Expi293 cells using ExpiFectamine293™ transfection reagents (Thermo Scientific, Waltham, MA), following manufacturer’s instructions. Supernatants containing the secreted antibodies were harvested, clarified and antibody purified using protein A chromatography. The purified antibodies were buffer exchanged in PBS and stored at 4°C.

### Biolayer interferometry (BLI)

8C1 affinity was measured on an Octet RED96e Biolayer Interferometer (Sartorius, Goettingen, Germany) using the Data Acquisition 12.0 software. Recombinant OspC_A_ was biotinylated using EZ-link NHS-biotin kit (Thermo Fisher Scientific). Biotinylated OspC (3 µg/mL) in PBS containing 2% w/v BSA (buffer) was captured onto Octet SA (streptavidin) biosensors (Sartorius) for 5 min. After 3 min to equilibrate to baseline in buffer, sensors were then immersed in a two-fold dilution series of 8C1 mAb or Fab, starting at 100 nM, for 10 min. The sensors were then dipped into wells containing buffer for 30 min to allow for dissociation. The raw sensor data was loaded into the Data Analysis HT 12.0 software, grouped and fit using a 1:2 bivalent analyte model (mAb) or a 1:1 model (Fab).

### *B. burgdorferi* motility determinations by dark field microscopy

Detailed strain description and methods associated with *B. burgdorferi* motility assays are described elsewhere ^52^. Briefly, mid-log-phase cultures of a *B. burgdorferi* B31 derivative (GGW94) carrying an IPTG-inducible *rpoS* allele was treated with OspC mAbs (1-30 μg/mL) in the absence or presence of 20% human complement (Sigma-Aldrich) for 16 or 24 h. After which cultures were examined in a double-blind fashion by dark-field microscopy for motile spirochetes using a Trinocular DF microscope (AmScope) equipped with a camera with reduction lens (AmScope SKU: MU1603) and a 40x dry darkfield condenser (AmScope; DK-DRY200). Spirochetes were considered nonviable when complete loss of motility and refractivity was observed. Spirochetes were enumerated in at least 4 visual fields, and the percent viability was calculated as the ratio of live spirochetes (mean of 4 fields) in treated samples to spirochetes in the untreated control samples (mean of 4 fields). Polyclonal serum from *B. burgdorferi*-infected mice and mAb B5 were used as positive controls; naive serum and the PB10 isotype were used as negative controls. This experimental set up was conducted over the course of three independent sessions and data is plotted as the means for the three days of counting. Statistical analysis was determined using 2-way ANOVA with Dunnett’s multiple comparisons test in which experimental means (8C1, B5) were compared to control mean (PB10) for each antibody concentration.

### Microsphere immunoassay (MIA)

Recombinant OspC_A_, OspC_B_, and OspC_K_ were coupled to MagPlex-C microspheres (5 μg antigen / 1×10^6^ microspheres) via the xMAP Antibody Coupling Kit (Luminex Corporation, Austin, TX) as recommended by the manufacturer. To couple biotinylated peptides, MagPlex-Avidin microspheres were suspended in assay buffer (1 x PBS, 2% BSA, pH 7.4) with biotinylated peptides (5 μg / 1×10^6^ microspheres) and allowed to incubate in a tabletop rotator at room temperature for 30 mins. Avidin microspheres were washed three times using a magnetic separator and wash buffer (1 x PBS, 2% BSA, 0.02% TWEEN-20, 0.05% Sodium azide, pH 7.4), resuspended in assay buffer, and stored at 2 – 8 until use.

Coupled microspheres were diluted in assay buffer (1:50) and then added (50 μL) to black, clear-bottomed, non-binding, chimney 96-well plates (Greiner Bio-One, Monroe, North Carolina). For the initial screens of hybridoma supernatant, 50 μL (neat) was added to each well. For the alanine mutant scan assays, 8C1 was diluted to 5 μg/mL and Lyme disease positive human serum samples kindly provided by the Lyme Disease Biobank at Nuvance Health^®^ (Danbury, CT) were diluted 1:100 in assay buffer and then added (50 μL) to each well ^37^. Plates were incubated for 1 h in a tabletop shaker (600 rpm) at room temperature and then washed 3 times using a magnetic separator and wash buffer. For the hybridoma supernatant screens, goat anti-mouse IgG, Human-ads-PE (SouthernBiotech, Birmingham, Alabama) secondary antibody was diluted 1:500 in assay buffer and added (100 μL) to each well. For the alanine mutant scan assays, PE labeled goat anti-Human IgG Fc, eBioscience (Invitrogen, Carlsbad, California) was diluted 1:500 in assay buffer and added (100 μL) to each well. Secondaries were incubated at room temperature for 30 min in a tabletop shaker (600 rpm). Plates were washed as previously stated, resuspended in 100 μL of wash buffer, and analyzed using a FlexMap 3D (Luminex Corporation). Both assay and wash buffers were prepared by the Wadsworth Center’s Cell and Tissue Culture core facility.

### HDX-MS

Stock concentrations of OspC-A (8.5 µM) in PBS either alone or in a complex with a 2-fold excess of antibody were diluted into 90 µL of deuterated PBS buffer (20 mM phosphate, 150 mM NaCl, 0.02% sodium azide, 1 mM EDTA pH* 7.54, 85%D final) containing 0.2 nM bradykinin and incubated 3 seconds on ice, or either 3 seconds, 1 minute, 30 min, or 20 hours at 21°C. Each starting stock also included a mixture of imidazolium compounds to serve as exchange reference standards ^53^. At the desired time point the sample was rapidly mixed with an equal volume of ice cold 0.2% formic acid and 0.1% trifluoroacetic acid (TFA) for a final pH of 2.5. Samples were then immediately frozen on ethanol/dry ice and stored at −80°C until LC-MS analysis. Undeuterated samples were prepared the same way but with undeuterated buffer for each step.

Samples were thawed at 5°C for 8 min and injected using a custom LEAP robot integrated with an LC-MS system ^54^. The protein was first passed over a Nepenthesin II column (2.1 x 30 mm; AffiPro) at 400 µL/min for inline digestion with the protease column held at 20°C. Peptides were then trapped on a Waters XSelect CSH C18 trap cartridge column (2.1 x 5 mm 2.5 µm) and resolved over a CSH C18 column (1 x 50 mm 1.7 µm 130Å) using linear gradient of 5 to 35% B (A: 0.1% FA, 0.025% TFA, 5% ACN; B: ACN with 0.1% FA) over 10 min and analyzed on a Thermo Orbitrap Ascend mass spectrometer at a resolution setting of 120,000. A series of washes over the trap and pepsin columns was used between injections to minimize carry-over as described ^54^. Data dependent MS/MS acquisition was performed on an undeuterated sample using rapid CID and HCD scans and processed in Byonic (Protein Metrics) with a score cutoff of 150 to identify peptides. Deuterium incorporation was analyzed using HDExaminer v3 (Sierra Analytics) ^55^.

## Financial Disclosure Statement

This work was supported by the National Institute of Allergy and Infectious Diseases (NIAID), National Institutes of Health, Department of Health and Human Services, Contract No. 75N93019C00040 (PI/PD Mantis). HDX instrumentation at the University of Washington was supported by award S10OD030237 from the National Institute of General Medical Sciences (NIGMS). The content is solely the responsibility of the authors and does not necessarily represent the official views of the National Institutes of Health.

## Competing Interests

The authors declare no competing financial interest.

## Acknowledgements

We gratefully acknowledge Mrs. Elizabeth Cavosie for administrative assistance. We thank the Applied Genomic Technologies core for DNA sequencing services, the Immunology core for access to flow cytometer, and the Media and Tissue Culture core for bacterial media. We gratefully acknowledge Dr. John Martignetti and Lisa Arrigo of the Nuvance Health Lyme disease Biobank for generously providing the Lyme disease serum samples used in this study.

## Supporting Information

**S1 Fig.** 8C1 hybridoma supernatant is OspC Type A specific. A) Hybridoma supernatant recognizes rOspC Type A but not B and K in Luminex. B) Dot blot analysis where 1×10^7^ bacteria or 1 µg of recombinant OspC protein types A, B, and K, were diluted and spotted on a nitrocellulose membrane and incubated with mouse hybridoma supernatant containing 8C1 IgG. The membrane was probed with an HRP-labelled anti-mouse IgG secondary antibody for detection of IgG binding. PBS and an unrelated protein, OspA were diluted and spotted as controls.

**S2 Fig.** Characterization of recombinant purified 8C1. A) Recombinant OspC Types A, B and K were coupled to Luminex beads, and then incubated with a dilution series of purified humanized 8C1. 8C1 displayed a clear dose response binding curve against rOspC_A_ but did not recognize OspC_B_ or OspC_K_ at any concentration tested. B-C) Biolayer interferometry sensorgrams of 8C1 mAb (B) or Fab (C) binding to OspC_A_.

**S3 Fig.** 8C1 displays complement-independent motility arrest. mAbs 8C1 (A), B5 (B) and isotype control mAb PB10 (C) were diluted serially and tested for their ability to inhibit *B. burgdorferi* mobility in a dark-field microscopy assay, with or without the addition of 20% human complement. A) At higher concentrations, 8C1 was able to inhibit more than 50% of spirochetal mobility in the absence of complement, an activity that was not statistically enhanced by the addition of complement. B) B5 displayed robust ability to arrest spirochete mobility in the absence of complement, and enhanced ability in the presence of complement. C) Isotype control mAb PB10 showed no dose dependent effect on sprirochete mobility with or without complement.

**S4 Fig.** 8C1 Fabs binding to OspC_A_-derived peptide 130-150. Biolayer interferometry sensorgrams of Fabs of 8C1 binding to a peptide composed of residues 130-150 from OspC_A_. The data is fit with a 1:1 binding model and gives a K_D_ of 9.2 nM.

## Notes

### Competing Interest Statement

The authors have declared no competing interest.

## References Cited

1. Steere, A.C., Strle, F., Wormser, G.P., Hu, L.T., Branda, J.A., Hovius, J.W., Li, X., and Mead, P.S. (2016). Lyme borreliosis. Nat Rev Dis Primers 2, 16090. 10.1038/nrdp.2016.90.

2. Radolf, J.D., Strle, K., Lemieux, J.E., and Strle, F. (2021). Lyme Disease in Humans. Curr Issues Mol Biol 42, 333–384. 10.21775/cimb.042.333.

3. Blum, L.K., Adamska, J.Z., Martin, D.S., Rebman, A.W., Elliott, S.E., Cao, R.R.L., Embers, M.E., Aucott, J.N., Soloski, M.J., and Robinson, W.H. (2018). Robust B Cell Responses Predict Rapid Resolution of Lyme Disease. Front Immunol 9, 1634. 10.3389/fimmu.2018.01634.

4. Bockenstedt, L.K., Hodzic, E., Feng, S., Bourrel, K.W., de Silva, A., Montgomery, R.R., Fikrig, E., Radolf, J.D., and Barthold, S.W. (1997). Borrelia burgdorferi strain-specific Osp C-mediated immunity in mice. Infect Immun 65, 4661–4667. 10.1128/iai.65.11.4661-4667.1997.

5. Barthold, S.W. (1999). Specificity of infection-induced immunity among Borrelia burgdorferi sensu lato species. Infect Immun 67, 36–42. 10.1128/IAI.67.1.36-42.1999.

6. Bhatia, B., Hillman, C., Carracoi, V., Cheff, B.N., Tilly, K., and Rosa, P.A. (2018). Infection history of the blood-meal host dictates pathogenic potential of the Lyme disease spirochete within the feeding tick vector. PLoS Pathog 14, e1006959. 10.1371/journal.ppat.1006959.

7. Barbour, A.G., and Travinsky, B. (2010). Evolution and distribution of the ospC Gene, a transferable serotype determinant of Borrelia burgdorferi. MBio 1. 10.1128/mBio.00153-10.

8. Khatchikian, C.E., Nadelman, R.B., Nowakowski, J., Schwartz, I., Wormser, G.P., and Brisson, D. (2014). Evidence for strain-specific immunity in patients treated for early lyme disease. Infect Immun 82, 1408–1413. 10.1128/IAI.01451-13.

9. Kurtti, T.J., Munderloh, U.G., Hughes, C.A., Engstrom, S.M., and Johnson, R.C. (1996). Resistance to tick-borne spirochete challenge induced by Borrelia burgdorferi strains that differ in expression of outer surface proteins. Infect Immun 64, 4148–4153. 10.1128/iai.64.10.4148-4153.1996.

10. Izac, J.R., and Marconi, R.T. (2019). Diversity of the Lyme Disease Spirochetes and its Influence on Immune Responses to Infection and Vaccination. Vet Clin North Am Small Anim Pract 49, 671–686. 10.1016/j.cvsm.2019.02.007.

11. Nadelman, R.B., Hanincova, K., Mukherjee, P., Liveris, D., Nowakowski, J., McKenna, D., Brisson, D., Cooper, D., Bittker, S., Madison, G., et al. (2012). Differentiation of reinfection from relapse in recurrent Lyme disease. N Engl J Med 367, 1883–1890. 10.1056/NEJMoa1114362.

12. De Silva, A.M., and Fikrig, E. (1995). Growth and migration of Borrelia burgdorferi in Ixodes ticks during blood feeding. Am J Trop Med Hyg 53, 397–404. 10.4269/ajtmh.1995.53.397.

13. Ohnishi, J., Piesman, J., and de Silva, A.M. (2001). Antigenic and genetic heterogeneity of Borrelia burgdorferi populations transmitted by ticks. Proc Natl Acad Sci U S A 98, 670–675. 10.1073/pnas.98.2.670.

14. Pal, U., Yang, X., Chen, M., Bockenstedt, L.K., Anderson, J.F., Flavell, R.A., Norgard, M.V., and Fikrig, E. (2004). OspC facilitates Borrelia burgdorferi invasion of Ixodes scapularis salivary glands. J Clin Invest 113, 220–230. 10.1172/JCI19894.

15. Schwan, T.G., Piesman, J., Golde, W.T., Dolan, M.C., and Rosa, P.A. (1995). Induction of an outer surface protein on Borrelia burgdorferi during tick feeding. Proc Natl Acad Sci U S A 92, 2909–2913. 10.1073/pnas.92.7.2909.

16. Eicken, C., Sharma, V., Klabunde, T., Owens, R.T., Pikas, D.S., Hook, M., and Sacchettini, J.C. (2001). Crystal structure of Lyme disease antigen outer surface protein C from Borrelia burgdorferi. J Biol Chem 276, 10010–10015. 10.1074/jbc.M010062200.

17. Kumaran, D., Eswaramoorthy, S., Luft, B.J., Koide, S., Dunn, J.J., Lawson, C.L., and Swaminathan, S. (2001). Crystal structure of outer surface protein C (OspC) from the Lyme disease spirochete, Borrelia burgdorferi. EMBO J 20, 971–978. 10.1093/emboj/20.5.971.

18. Zuckert, W.R., Kerentseva, T.A., Lawson, C.L., and Barbour, A.G. (2001). Structural conservation of neurotropism-associated VspA within the variable Borrelia Vsp-OspC lipoprotein family. J Biol Chem 276, 457–463. 10.1074/jbc.M008449200.

19. Caimano, M.J., Groshong, A.M., Belperron, A., Mao, J., Hawley, K.L., Luthra, A., Graham, D.E., Earnhart, C.G., Marconi, R.T., Bockenstedt, L.K., et al. (2019). The RpoS Gatekeeper in Borrelia burgdorferi: An Invariant Regulatory Scheme That Promotes Spirochete Persistence in Reservoir Hosts and Niche Diversity. Front Microbiol 10, 1923. 10.3389/fmicb.2019.01923.

20. Barbour, A.G., Jasinskas, A., Kayala, M.A., Davies, D.H., Steere, A.C., Baldi, P., and Felgner, P.L. (2008). A genome-wide proteome array reveals a limited set of immunogens in natural infections of humans and white-footed mice with Borrelia burgdorferi. Infect Immun 76, 3374–3389. 10.1128/IAI.00048-08.

21. Di, L., Wan, Z., Akther, S., Ying, C., Larracuente, A., Li, L., Di, C., Nunez, R., Cucura, D.M., Goddard, N.L., et al. (2018). Genotyping and Quantifying Lyme Pathogen Strains by Deep Sequencing of the Outer Surface Protein C (ospC) Locus. J Clin Microbiol 56. 10.1128/JCM.00940-18.

22. Baum, E., Randall, A.Z., Zeller, M., and Barbour, A.G. (2013). Inferring epitopes of a polymorphic antigen amidst broadly cross-reactive antibodies using protein microarrays: a study of OspC proteins of Borrelia burgdorferi. PLoS One 8, e67445. 10.1371/journal.pone.0067445.

23. Theisen, M., Borre, M., Mathiesen, M.J., Mikkelsen, B., Lebech, A.M., and Hansen, K. (1995). Evolution of the Borrelia burgdorferi outer surface protein OspC. J Bacteriol 177, 3036–3044. 10.1128/jb.177.11.3036-3044.1995.

24. Earnhart, C.G., Buckles, E.L., Dumler, J.S., and Marconi, R.T. (2005). Demonstration of OspC type diversity in invasive human lyme disease isolates and identification of previously uncharacterized epitopes that define the specificity of the OspC murine antibody response. Infect Immun 73, 7869–7877. 10.1128/IAI.73.12.7869-7877.2005.

25. Wilske, B., Preac-Mursic, V., Jauris, S., Hofmann, A., Pradel, I., Soutschek, E., Schwab, E., Will, G., and Wanner, G. (1993). Immunological and molecular polymorphisms of OspC, an immunodominant major outer surface protein of Borrelia burgdorferi. Infect Immun 61, 2182–2191.

26. Probert, W.S., Crawford, M., Cadiz, R.B., and LeFebvre, R.B. (1997). Immunization with outer surface protein (Osp) A, but not OspC, provides cross-protection of mice challenged with North American isolates of Borrelia burgdorferi. J Infect Dis 175, 400–405.

27. Gomes-Solecki, M., Arnaboldi, P.M., Backenson, P.B., Benach, J.L., Cooper, C.L., Dattwyler, R.J., Diuk-Wasser, M., Fikrig, E., Hovius, J.W., Laegreid, W., et al. (2020). Protective Immunity and New Vaccines for Lyme Disease. Clin Infect Dis 70, 1768–1773. 10.1093/cid/ciz872.

28. Burton, D.R. (2019). Advancing an HIV vaccine; advancing vaccinology. Nat Rev Immunol 19, 77–78. 10.1038/s41577-018-0103-6.

29. Angeletti, D., and Yewdell, J.W. (2018). Understanding and Manipulating Viral Immunity: Antibody Immunodominance Enters Center Stage. Trends Immunol 39, 549–561. 10.1016/j.it.2018.04.008.

30. Caradonna, T.M., and Schmidt, A.G. (2021). Protein engineering strategies for rational immunogen design. NPJ Vaccines 6, 154. 10.1038/s41541-021-00417-1.

31. Mittal, A., Khattri, A., and Verma, V. (2022). Structural and antigenic variations in the spike protein of emerging SARS-CoV-2 variants. PLoS Pathog 18, e1010260. 10.1371/journal.ppat.1010260.

32. Rudolph, M.J., Davis, S.A., Haque, H.M.E., Weis, D.D., Vance, D.J., Piazza, C.L., Ejemel, M., Cavacini, L., Wang, Y., Mbow, M.L., et al. (2023). Structural Elucidation of a Protective B Cell Epitope on Outer Surface Protein C (OspC) of the Lyme Disease Spirochete, Borreliella burgdorferi. mBio 14, e0298122. 10.1128/mbio.02981-22.

33. Rudolph, M.J., Chen, Y., Vorauer, C., Vance, D.J., Piazza, C.L., Willsey, G.G., McCarthy, K., Muriuki, B., Cavacini, L.A., Guttman, M., and Mantis, N.J. (2024). Structure of a Human Monoclonal Antibody in Complex with Outer Surface Protein C of the Lyme Disease Spirochete, Borreliella burgdorferi. J Immunol. 10.4049/jimmunol.2400247.

34. Hyde, J.A. (2017). Borrelia burgdorferi Keeps Moving and Carries on: A Review of Borrelial Dissemination and Invasion. Front Immunol 8, 114. 10.3389/fimmu.2017.00114.

35. Dang, X., Guelen, L., Lutje Hulsik, D., Ermakov, G., Hsieh, E.J., Kreijtz, J., Stammen-Vogelzangs, J., Lodewijks, I., Bertens, A., Bramer, A., et al. (2023). Epitope mapping of monoclonal antibodies: a comprehensive comparison of different technologies. MAbs 15, 2285285. 10.1080/19420862.2023.2285285.

36. Buckles, E.L., Earnhart, C.G., and Marconi, R.T. (2006). Analysis of antibody response in humans to the type A OspC loop 5 domain and assessment of the potential utility of the loop 5 epitope in Lyme disease vaccine development. Clin Vaccine Immunol 13, 1162–1165. 10.1128/CVI.00099-06.

37. Freeman-Gallant, G., McCarthy, K., Yates, J., Kulas, K., Rudolph, M.J., Vance, D.J., and Mantis, N.J. (2024). A refined human linear B cell epitope map of Outer surface protein C (OspC) from the Lyme disease spirochete, Borreliella burgdorferi. bioRxiv, 2024.2005.2029.596441. 10.1101/2024.05.29.596441.

38. Yang, X., Li, Y., Dunn, J.J., and Luft, B.J. (2006). Characterization of a unique borreliacidal epitope on the outer surface protein C of Borrelia burgdorferi. FEMS Immunol Med Microbiol 48, 64–74. 10.1111/j.1574-695X.2006.00122.x.

39. Earnhart, C.G., and Marconi, R.T. (2007). Construction and analysis of variants of a polyvalent Lyme disease vaccine: approaches for improving the immune response to chimeric vaccinogens. Vaccine 25, 3419–3427. 10.1016/j.vaccine.2006.12.051.

40. Tokarz, R., Mishra, N., Tagliafierro, T., Sameroff, S., Caciula, A., Chauhan, L., Patel, J., Sullivan, E., Gucwa, A., Fallon, B., et al. (2018). A multiplex serologic platform for diagnosis of tick-borne diseases. Sci Rep 8, 3158. 10.1038/s41598-018-21349-2.

41. Shandilya, S., Kurt Yilmaz, N., Sadowski, A., Monir, E., Schiller, Z.A., Thomas, W.D., Jr., Klempner, M.S., Schiffer, C.A., and Wang, Y. (2017). Structural and molecular analysis of a protective epitope of Lyme disease antigen OspA and antibody interactions. J Mol Recognit 30. 10.1002/jmr.2595.

42. Yamashita, T., Mizohata, E., Nagatoishi, S., Watanabe, T., Nakakido, M., Iwanari, H., Mochizuki, Y., Nakayama, T., Kado, Y., Yokota, Y., et al. (2019). Affinity Improvement of a Cancer-Targeted Antibody through Alanine-Induced Adjustment of Antigen-Antibody Interface. Structure 27, 519–527 e515. 10.1016/j.str.2018.11.002.

43. Fraser, C.M., Casjens, S., Huang, W.M., Sutton, G.G., Clayton, R., Lathigra, R., White, O., Ketchum, K.A., Dodson, R., Hickey, E.K., et al. (1997). Genomic sequence of a Lyme disease spirochaete, Borrelia burgdorferi. Nature 390, 580–586. 10.1038/37551.

44. Anderson, J.F., Flavell, R.A., Magnarelli, L.A., Barthold, S.W., Kantor, F.S., Wallich, R., Persing, D.H., Mathiesen, D., and Fikrig, E. (1996). Novel Borrelia burgdorferi isolates from Ixodes scapularis and Ixodes dentatus ticks feeding on humans. J Clin Microbiol 34, 524–529. 10.1128/jcm.34.3.524-529.1996.

45. Steere, A.C., Grodzicki, R.L., Kornblatt, A.N., Craft, J.E., Barbour, A.G., Burgdorfer, W., Schmid, G.P., Johnson, E., and Malawista, S.E. (1983). The spirochetal etiology of Lyme disease. N Engl J Med 308, 733–740. 10.1056/NEJM198303313081301.

46. Barbour, A.G. (1984). Isolation and cultivation of Lyme disease spirochetes. Yale J Biol Med 57, 521–525.

47. Rudolph, M.J., Davis, S.A., Haque, H.M.E., Ejemel, M., Cavacini, L.A., Vance, D.J., Willsey, G.G., Piazza, C.L., Weis, D.D., Wang, Y., and Mantis, N.J. (2023). Structure of a transmission blocking antibody in complex with Outer surface protein A from the Lyme disease spirochete, Borreliella burgdorferi. Proteins. 10.1002/prot.26549.

48. Frye, A.M., Ejemel, M., Cavacini, L., Wang, Y., Rudolph, M.J., Song, R., and Mantis, N.J. (2022). Agglutination of Borreliella burgdorferi by Transmission-Blocking OspA Monoclonal Antibodies and Monovalent Fab Fragments. Infect Immun, e0030622. 10.1128/iai.00306-22.

49. Harlow, E., and Lane, D. (1988). Antibodies: A laboratory manual. (Cold Spring Harbor Laboratory Press).

50. Van Slyke, G., Angalakurthi, S.K., Toth, R.T.t., Vance, D.J., Rong, Y., Ehrbar, D., Shi, Y., Middaugh, C.R., Volkin, D.B., Weis, D.D., and Mantis, N.J. (2018). Fine-Specificity Epitope Analysis Identifies Contact Points on Ricin Toxin Recognized by Protective Monoclonal Antibodies. Immunohorizons 2, 262-273. 10.4049/immunohorizons.1800042.

51. Meyer, L., Lopez, T., Espinosa, R., Arias, C.F., Vollmers, C., and DuBois, R.M. (2019). A simplified workflow for monoclonal antibody sequencing. PLoS One 14, e0218717. 10.1371/journal.pone.0218717.

52. Rudolph, M.J., Chen, Y., Vorauer, C., Vance, D.J., Piazza, C.L., Willsey, G.G., McCarthy, K., Muriuki, B., Cavacini, L.A., Guttman, M., and Mantis, N.J. (2024). Structure of a human monoclonal antibody in complex with Outer surface protein C (OspC) of the Lyme disease spirochete, Borreliella burgdorferi. bioRxiv. 10.1101/2024.04.29.591597.

53. Murphree, T.A., Vorauer, C., Brzoska, M., and Guttman, M. (2020). Imidazolium Compounds as Internal Exchange Reporters for Hydrogen/Deuterium Exchange by Mass Spectrometry. Analytical Chemistry 92, 9830–9837. 10.1021/acs.analchem.0c01328.

54. Watson, M.J., Harkewicz, R., Hodge, E.A., Vorauer, C., Palmer, J., Lee, K.K., and Guttman, M. (2021). Simple Platform for Automating Decoupled LC-MS Analysis of Hydrogen/Deuterium Exchange Samples. J Am Soc Mass Spectrom 32, 597–600. 10.1021/jasms.0c00341.

55. Hageman, T.S., and Weis, D.D. (2019). Reliable Identification of Significant Differences in Differential Hydrogen Exchange-Mass Spectrometry Measurements Using a Hybrid Significance Testing Approach. Anal Chem 91, 8008–8016. 10.1021/acs.analchem.9b01325.

