## Supplementary figures and images for "A type-specific B cell epitope at the apex of Outer surface protein C (OspC) of the Lyme disease spirochete, *Borreliella burgdorferi*"

### Supplemental Figure 1-4

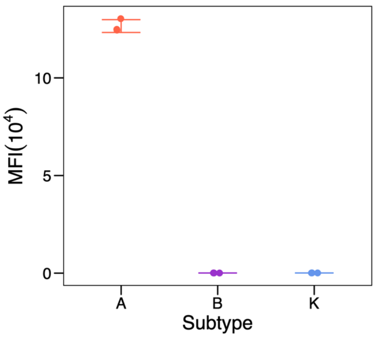

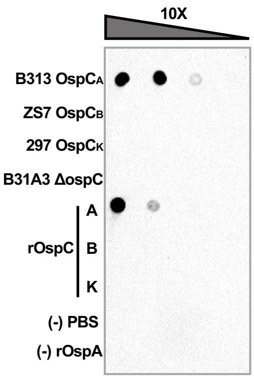

**Figure S2**

**A.**

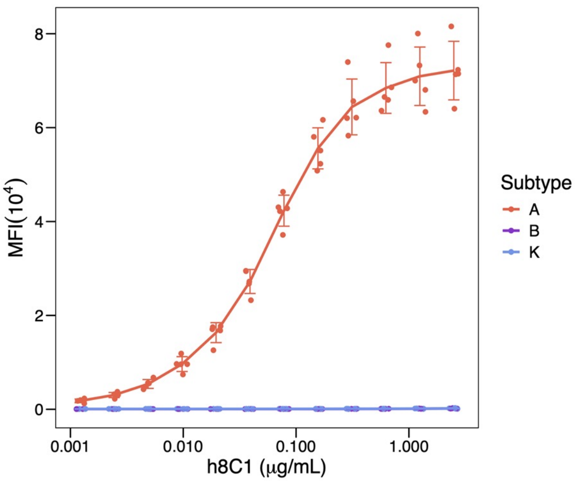


**B.**

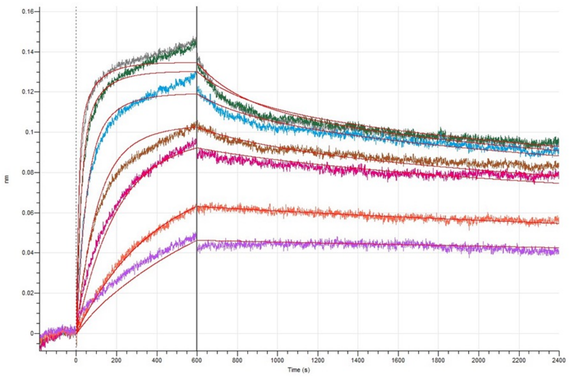


**C.**

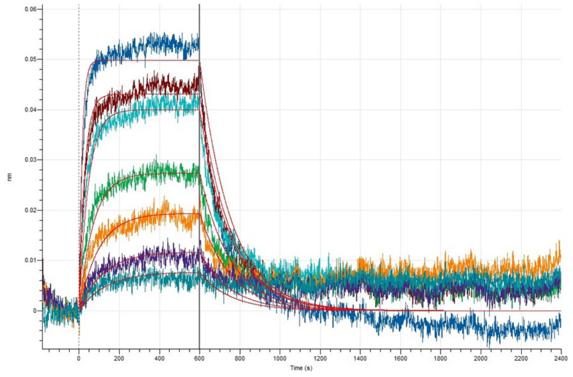

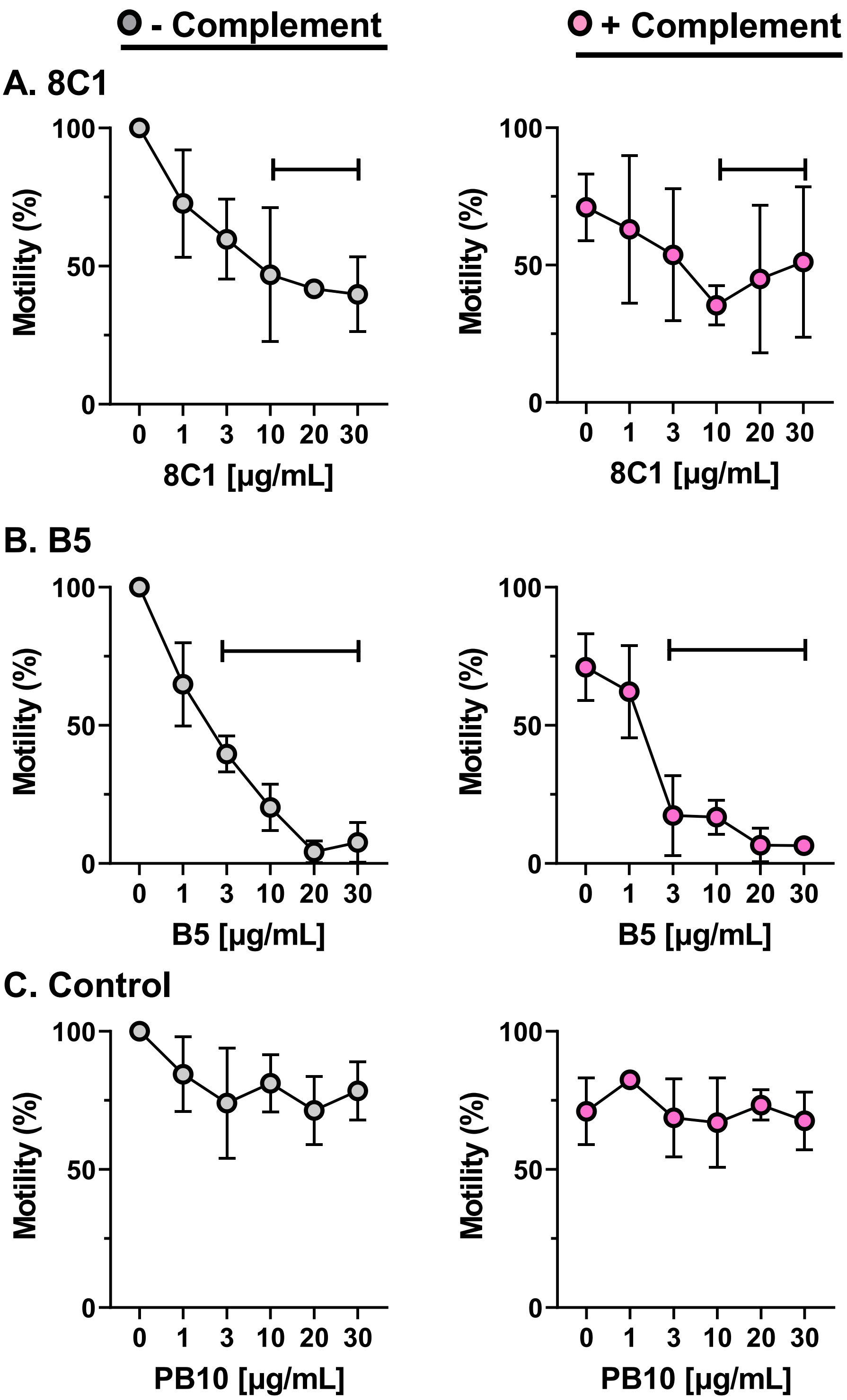

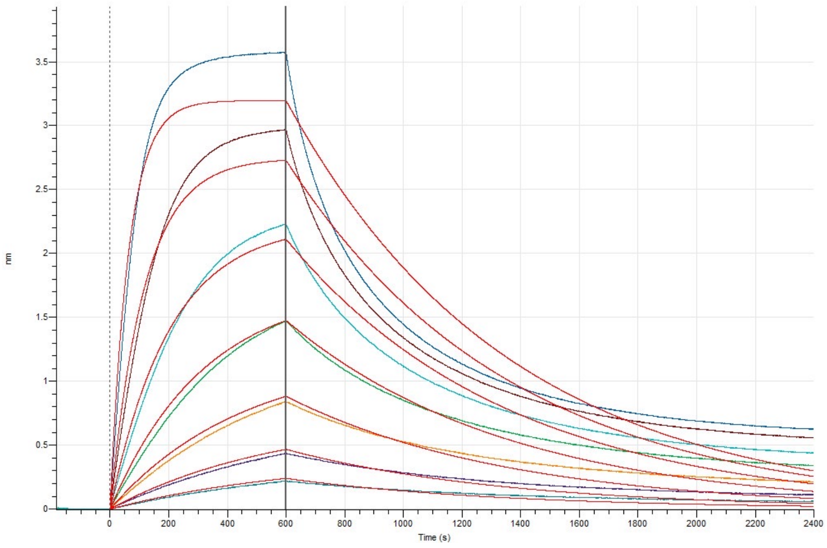
